# Beyond Structural Bioinformatics for Genomics with Dynamics Characterization of an Expanded KRAS Mutational Landscape

**DOI:** 10.1101/2023.04.28.536249

**Authors:** Brian D. Ratnasinghe, Neshatul Haque, Jessica B. Wagenknecht, Davin R. Jensen, Guadalupe V. Esparza, Elise N. Leverence, Thiago Milech De Assuncao, Angela J. Mathison, Gwen Lomberk, Brian C. Smith, Brian F. Volkman, Raul Urrutia, Michael T. Zimmermann

## Abstract

Current capabilities in genomic sequencing outpace functional interpretations. Our previous work showed that 3D protein structure calculations enhance mechanistic understanding of genetic variation in sequenced tumors and patients with rare diseases. The KRAS GTPase is among the critical genetic factors driving cancer and germline conditions. Because KRAS-altered tumors frequently harbor one of three classic hotspot mutations, nearly all studies have focused on these mutations, leaving significant functional ambiguity across the broader KRAS genomic landscape observed in cancer and non-cancer diseases. Herein, we extend structural bioinformatics with molecular simulations to study an expanded landscape of 86 KRAS mutations. We identify multiple coordinated changes strongly associated with experimentally established KRAS biophysical and biochemical properties. The patterns we observe span hotspot and non-hotspot alterations, which can all dysregulate Switch regions, producing mutation-restricted conformations with different effector binding propensities. We experimentally measured mutation thermostability and identified shared and distinct patterns with simulations. Our results indicate mutation-specific conformations which show potential for future research into how these alterations reverberate into different molecular and cellular functions. The data we present is not predictable using current genomic tools, demonstrating the added functional information derived from molecular simulations for interpreting human genetic variation.

**Key Points:** Please provide 3 bullet points summarizing the manuscript’s contribution to the field (100 characters max per point)

1) We functionally grouped 86 distinct KRAS mutations using MD scores, demonstrating scalability for genomics
2) MD-based groups explain experimental differences and mechanistic information about mutant proteins
3) Demonstrated added functional information from simulations for interpreting human genetic variation

## INTRODUCTION

A significant unmet need is determining which uninterpreted genetic variants (variants of unknown significance, VUS) carry biomedical significance, such as in proto-oncogenes. Currently, variants are classified using guidelines that mainly rely on genetic observations and linear models of DNA and protein sequences(1,2). Appropriate incorporation of mechanistic biophysical and biochemical features of the encoded 3D molecule will significantly improve the accuracy of interpreting disease-associated genomic variation, compared to existing genomics tools. Current genomics tools predict missense mutations in the proto-oncogene KRAS (the rat sarcoma viral oncogene homolog discovered by Kirsten) to be uniformly damaging. Yet, experiments show that different mutations have distinct properties. KRAS enzymatic activity is conformationally regulated by specific loops called the Switch regions, whose positions also determine whether KRAS binds to other effector proteins. Switch conformations differ between the GDP and GTP-bound states, such that the GTP-bound form binds the effectors that activate the downstream cell signaling cascade. Thus, enzymatic and cell signaling activities interchange, with enzyme activity turning off cellular activity, while compromised GTPase activity leads to greater cellular activity. A previous study of KRAS G12D compared to G13D (3), for example, demonstrated that mutations lead to alternate Switch positions that define mutation-specific conformations and change binding to effector proteins. However, only the most common mutations have been characterized in depth, leaving significant ambiguity about the intrinsic effects of the expanded mutational landscape. Thus, KRAS is an excellent model system to advance biomedical knowledge and develop specific methodologies for using molecular simulations to interpret inter-individual genetic variation.

The current study illuminates specific details about KRAS mutant proteins. We recently used experimental structures across the RAS GTPase family to increase the possible scale, leveraging structure bioinformatics to interpret human genetic variation (4,5). Previously, the most extensive KRAS study of this type analyzed six different G12 position variants (6). Herein, we use physics-based simulations and experimental biophysical validation to bring new and uniform information across a broad landscape of 86 KRAS mutations observed in cancer and non-cancer diseases. Our simulations capture dynamic movements across the protein, with the largest movements observed for the Switch regions. We demonstrate that each mutation produces shared and distinct protein structural and dynamic features. Differences across intrinsic features of KRAS mutant proteins will set the stage for how the enzyme behaves within human cells and across body tissues, shaping cellular responses. The data generated and knowledge derived will therefore transfer to different contexts, such as across tumors from other body tissues, empowering the interpretation of existing and future studies.

## MATERIAL AND METHODS

### Developing and Generating Molecular Dynamics Simulation Data

The initial conformations were taken from Chain A of the X-ray crystallography structure from the Protein Data Bank (7) of human WT KRAS bound to GDP-Mg^2+^ (4OBE (8)). FoldX v4 (9) was used for computational mutagenesis, generating 86 variant structures. The 86 variants were selected according to incidence across human cancers and inherited germline RASopathies, concordant with our previous studies(5). Each mutant KRAS structure was modeled bound to GTP-Mg^2+^ by fitting the 6OB2 (10) GDP-PnP to GDP and converting it into GTP. After these initial steps, we have two versions of each mutation, plus WT, wherein one has GTP bound, and the other has GDP bound.

All structures were prepared for simulations with a uniform initial environment. The CHARMM36 force field (11–14), CHARMM-modified TIP3P water (15–17) (including Lennard-Jones terms of hydrogen bonds), and standard CHARMM ion parameters were used (18). Our KRAS-WT model was centered in a cubic cell with a box-solute minimum distance of 10 Å, filled with TIP3P and neutralizing counter-ions up to 150 mM KCl. Each mutant KRAS model was placed in this initial environment.

Each system was independently energy-minimized using NAMD(19) with 10,000 steps of steepest decent minimization. Three independent replicates were produced for each KRAS mutation by generating different random velocities at the outset of equilibration. All protein and ligand atoms were restrained to their initial positions using a 10kcal mol^-1^ Å^-2^ harmonic constraint. The energy was initialized at 10K and following the Boltzmann distribution. The energy was slowly increased to 300K over 0.7ns with position restraints applied to all nonhydrogen protein atoms. Then, restraints were released; initially lowered to 5kcal mol^-1^ Å^-2^ for 0.1ns, then 2kcal mol^-1^ Å^-2^ for 0.1ns, and gradually released by increments of 0.05kcal mol^-1^ Å^-2^ for 20ps each until they reached zero. Then free equilibration was performed for 12ns under the NPT ensemble, implemented via the Langevin thermostat method at 300K with a friction coefficient of 5ps^-1^. Water molecules and ions were allowed to diffuse freely during equilibration time. The Langevin piston method (20) was applied to keep the pressure at 1 atm with an oscillation period of 400fs, a decay time of 100fs, and an isotopically scaled box. Periodic Boundary Conditions were applied in all three spatial directions. The particle mesh Ewald method (21,22) was used to calculate electrostatic interactions with a real-space cutoff of 12 Å and a grid spacing of ∼1Å. Then the short-range van der Waals forces were switched from 10 to 12 Å. All bonds involving hydrogen atoms were constrained using the SHAKE algorithm (23), and all water molecules were constrained with SETTLE (24), with an integration timestep of 1fs.

After these system preparations, NAMD was also used to perform unrestrained simulations that extended from each equilibration run. These were performed in NVT ensemble for 40ns each and a total of 120ns for each variant in both GDP and GTP bound forms. The full simulation time between all variants and nucleotides was 21μs. The temperature in production simulations was maintained using the Anderson thermostat (25) with a 1 ps^-1^ collision frequency. The pressure was isotopically regulated with the Monte-Carlo barostat, with box scaling attempted every 20 integration steps. Other parameters were the same as during equilibration.

### Structural Calculations and Statistical Comparisons

The root-mean-square deviation (RMSD) and root-mean-square fluctuation (RMSF) values were calculated using C^α^ atoms after structurally aligning each trajectory to the initial WT conformation and ignoring the two mobile Switch loops. Principal Component (PC) analysis summarized the dominant conformational changes across trajectories. PCs were calculated in Cartesian coordinates and using C^α^ atoms. An observed Free Energy Landscape (FEL) was calculated for each PC-based subspace using the approach of Karamzadeh *et al*. (26) Differences between each mutation and WT were quantified using a distributional comparison. In this approach, the sampling of PC1 by a mutation was scored as the number of standard deviations in its mean, from the WT-mean and WT-variation along PC1. This procedure was done for each of the top 3 PCs, providing standardized scores for each mutation.

Amino acid pairwise distances used as markers for key experimentally determined biophysical or biochemical properties of KRAS and other RAS-family proto-oncogenes from the literature (3,27,28). Given the robust data supporting each, they were used to interpret our dynamics-based data. When stretching or compressing the distances between the amino acid pairs was used, for example, the changes observed in the simulation were interpreted according to the biophysically and biochemically derived precedent. Thermostability measurements (described below) were also used for validation, which in this study have the unprecedented advantage, over previous studies, of being uniformly generated across the mutational landscape. Significance was scored according to how many standard deviations (denoted with σ) from WT-median values, each mutation’s median was observed. Multiple structure- and dynamics-based scores and calculations were combined, generating cross-correlation matrices to understand their inter-relationships and UMAP (29) visualizations for the relative similarities of KRAS mutations across the dataset.

Analyses were performed out using a custom structural bioinformatics workflow and leveraging the bio3d R package (30) Protein annotations were gathered from UniProt (31) and the literature. Electrostatics were calculated using APBS (32) Protein structures and simulation trajectories were visualized using VMD (33) version 1.9.3 and PyMOL (34) versions 2.3.0 and 2.6.0a0.

### Establishing Meta Classifications of KRAS Mutant Proteins

Multiple calculations were made, and each was used to classify mutations based on their similarity to WT. The GTP and GDP interaction energy scores were split into nine categories: Variants within one standard deviation (1σ) of the WT mean are considered neutral variants; variants within 1σ of WT GDP but more than 1σ away from WT GTP are referred to as “GDP Stable” or “GTP Unstable”. Conversely, variants within 1σ from WT GTP but more than 1σ away from WT GDP are referred to as “GTP Stable” or “GDP Unstable”. Variants that do not have GDP or GTP values within 1σ of WT will either be overly stable or unstable for both nucleotides. As such, variants were categorized as Stable, Neutral, and Unstable in two dimensions (GDP and GTP energy) to produce 9 groups of variants. RMSD scores, calculated from RAS NF1 (PDB: 6OB2) and SOS1 (PDB: 6EPM) bound conformations, were split according to WT-like or deviated, according to their relative difference from WT in both GDP and GTP states. To score changes in dynamics sampling as measured by PC analysis, Interaction Energy, RMSD and distance monitor sampling, we split variants into three categories each – those that exhibited significantly lower, WT-like, or significantly higher values; variants ≥1σ from the mean classified as high, while variants ≤-1σ are classified as low. To score changes in dynamics sampling as measured by PC analysis, RMSD and distance monitor sampling, we split variants into three categories each – those that exhibited significantly lower, WT-like, or significantly higher values; variants ≥1σ from the mean classified as high, while variants ≤-1σ are classified as low. This procedure was applied to 20 scores, producing 108 meta-classifications. Of these 108 meta-classifications we filtered those that were not among the top 10 defining features of any k-means groups to generate a signature for each k-means group across the many meta- classifications to account for multiple protein features concurrently. In total, twelve K-means clustering groups of meta-class signatures were generated, with six groups in each GTP and GDP state. Analyses were conducted using custom scripts in the R programming language (35). The entire matrix of scores across mutations was visualized using the pheatmap package (36).

### Expression and Purification of KRAS Mutant Proteins

The KRAS expression plasmid was obtained from Addgene (Plasmid #111849) as an E. Coli codon optimized plasmid encoding A.A. 1-169. Plasmids were transformed into E. Coli strain BL21(DE3) and grown at 37°C in 50mL of M9 media with ^15^N ammonium chloride to make uniformly ^15^N- labeled KRAS. Cells stayed at 37°C until the absorbance at 600nm was 0.7, then 1mM IPTG was added, and temperature adjusted to 18°C and induced overnight. The next day the cells were pelleted by centrifugation 5000xg and stored at -20°C until further processing.

KRAS pellets were resuspended in 50mM HEPES, 300mM NaCl, 10mM imidazole, 5% glycerol, 0.1% beta mercaptoethanol, 0.1mg/mL Lysozyme and 1mM MgCl_2_. Then the pellets were lysed by sonication, centrifuged 30 minutes at 15,000g, and the soluble fraction kept for nickel affinity purification. KRAS proteins were purified in workgroups of 12, using the Maxwell 16 (Promega) utilizing magnetic nickel beads in a semi-automated nickel affinity purification strategy. The Maxwell 16 has a cassette with 7 wells; the clarified cell lysate is placed in well 1, wash buffers (50mM HEPES, pH 7.5, 300mM NaCl, 10mM imidazole and 5% glycerol) in wells 2-6, and 150uL of magnetic resin also in well 2. The instrument will move the resin to well 1 to bind the protein of interest, wash the resin in wells 3-6 and finally elute the protein in a separate cuvette. After purification, elutions are dialyzed against 50mM HEPES, pH 7.0, 50mM NaCl, 10mM MgCl_2_ and 1mM Dtt.

### Nucleotide Exchange Experiments for Producing GDP and GTP-Analog-Bound Proteins

To prepare enough KRAS with the desired nucleotide, about 100uL of 100uM KRAS was placed into a spin concentrator. Then 300uL of ‘low magnesium’ buffer (10mM HEPES, pH 7.5, 50mM NaCl, 500uM MgCl_2_ and 2mM Dtt) was added and concentrated to 100uL, then the buffer addition and concentration to 100uL was repeated 2 more times. To exchange the nucleotide, 100uL of Nucleotide exchange buffer (20mM HEPES, pH 7.0, 50mM NaCl, 10mM EDTA, 20-fold molar excess concentration of nucleotide, GTP or GPPnP) was added and left on ice for 90 minutes. Then 20mM MgCl_2_ was added to ‘lock’ the nucleotide in place and left on ice for another 30 minutes before using a desalting column to exchange the protein into 10mM HEPES pH 7.0, 20mM NaCl, 5mM MgCl_2_.

### Measurement of KRAS Thermostabilities

The NanoTemper Prometheus (NanoTemper Technologies) is a nano-DSF that can determine Thermal unfolding temperatures using the intrinsic fluorescence of the protein over a temperature gradient. The instrument uses glass capillaries that only require 10uL of 1mg/mL for KRAS. After nucleotide exchange, samples of each protein in both the GDP and GPPnP state were diluted to 1mg/mL. Then a capillary is placed in the sample and filled with capillary action. Temperature gradients were 20-95°C at a speed of 0.5°C/min. The melting temperature is determined to be peak maximum of the first derivative of the 350/330nm fluorescence ratio.

## RESULTS

The intrinsic hydrolytic activity of KRAS is primarily regulated at a conformational level by two loops designated Switch 1 and Switch 2 (**Figure 1A**). Specific conformational changes, differently stabilized by GTP and GDP, coincide with binding different nucleotide exchange factors (GEFs), activating proteins (GAPs), or additional signaling proteins (effectors). We generate and analyze Molecular Dynamics (MD) to observe the different Switch conformations across an expanded KRAS mutational landscape. These calculations allow us to distinguish different Switch dynamics for each distinct mutation, and then group mutations by their similarity in dynamic behavior. Our goal is to better interpret the nuanced effects of each genetic variation on the encoded protein. Indeed, there is even potential to use these dynamics scores to develop a robust model capable of providing direct insight on how KRAS mutations cause disease, spanning classic hotspot, non-classic hotspot, and non-hotspot genetic variants.

**Figure 1.**
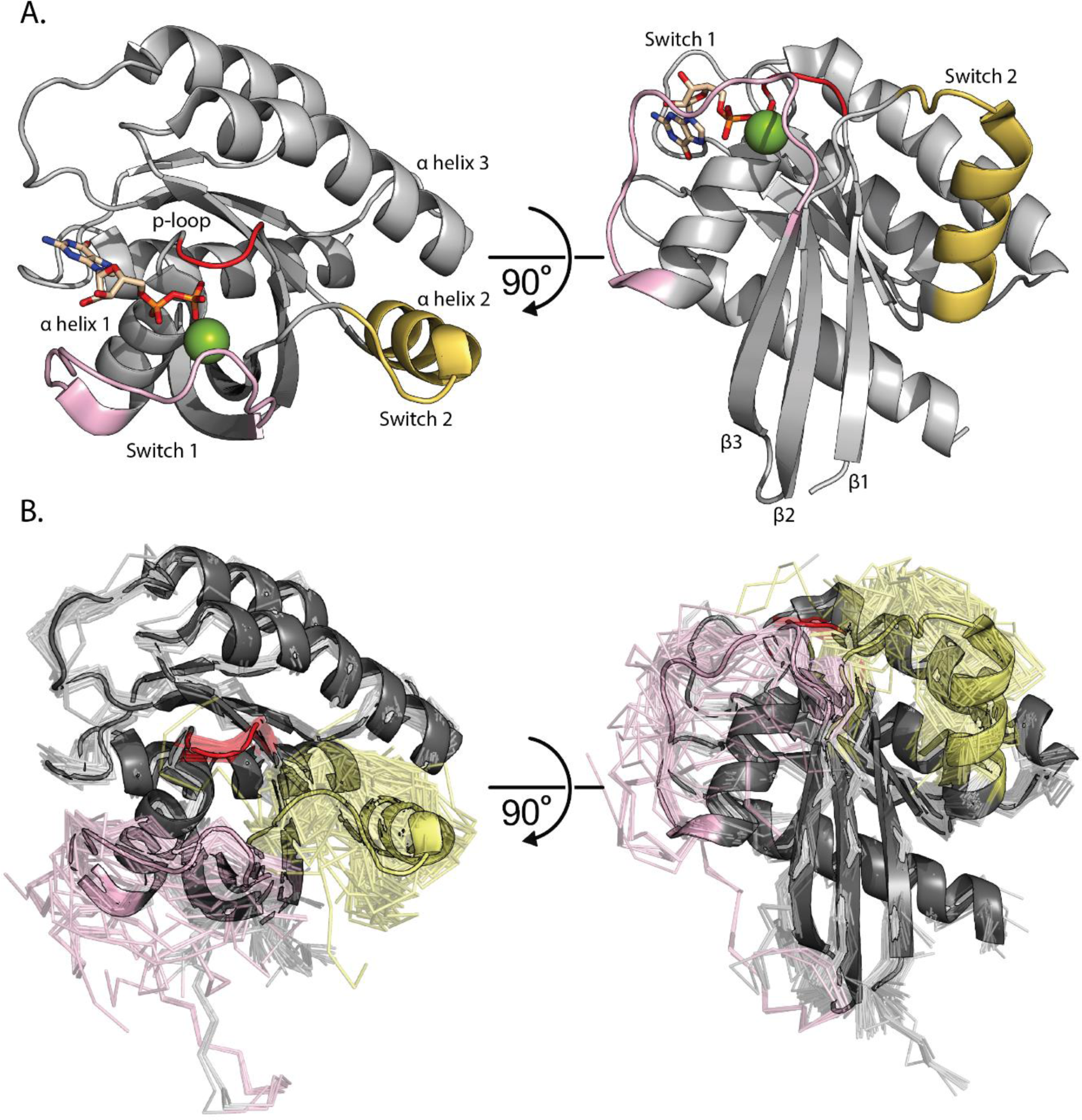
RAS Structural Overview. A) KRAS Structure colored in grey bound to GDP with carbons in tan and Mg^2+^ colored Green (PDB: 4OBE). The p-loop is colored red, Switch 1 is colored pink, and Switch 2 is colored yellow. Additionally, α helices 1, 2 and 3 are labeled where α helix 1 is the first helix in the KRAS sequence and α helix 3 is the third helix in the sequence. The same KRAS structure with the same coloring rotated 60°. Here, β sheets 1, 2, and 3 are labeled where β sheet 1 is the first sheet in the KRAS sequence and β sheet 3 is the third sheet in the sequence. B) Complete representation of all 194 PDB deposited KRAS structures (chain A). In the cartoon, GDP bound WT structure is used as the starting point for simulations (PDB: 4OBE). The other 193 KRAS conformations are displayed in ribbon. The p-loop is colored red, Switch 1 is colored pink, Switch 2 is colored yellow.

### Experimental KRAS Structural Characterization Has Focused on a Narrow Field of Mutations

We generated an experimental data corpus of KRAS Switch dynamics across multiple functional and ligand-bound states. Using this curated dataset, we quantified the current scope of KRAS structural studies and measured how extensive our simulations sampled the experimental landscape. We identified 194 experimentally solved KRAS structures (**Figure 1B**) and found that 73 (38%) do not have Switch 1 or Switch 2 fully resolved, meaning these loops were too dynamic in the protein crystal to assign coordinates. Therefore, current experimental data can be extended using simulations to enhance our understanding of Switch functional conformations and their mutational dependencies. Additionally, only 18 distinct mutants have experimentally solved structures, further demonstrating the utility of MD in providing unique, complementary, and highly detailed information on a large scale (**Table S1**). Of these 18 mutants, WT, G12C, and G12D comprise 137 (71%) of the 194 experimental structures (**Table S1**). Only 36 (19%) are with KRAS in complex with other proteins (detailed in **Table S1**), of which 22/36 (61%) of these complexes feature a nanodisc with an engineered RAS binding protein (DARPins and affirmers). Only 7/36 (19%) conformations are in complex with physiologic binding partners such as NF1 (GAP), SOS1 (GEF), and RAF1 (effector). In summary, there is a lack of uniformity across the experimentally determined ensemble of KRAS structures, demonstrating the need for a uniform and systematic approach to obtaining structural and dynamic information across a broad KRAS mutational landscape.

### Coordinate PC Landscape Differentiates Among Groups of Mutations

From our MD simulations, we uniformly and systematically quantified differences in conformational sampling among mutant KRAS proteins by first performing Principal Component (PC) analyses. We focused on the top 3 PCs, which capture the most dominant movements, collectively describing 37.7% of the conformational variation across all simulations. PC1 describes the overall dynamics of Switch 1 while PC2 and PC3 represent different Switch 2 dynamics (**Figure S1**). We used a free energy landscape approach to summarize the sampling of the top PCs from GTP (**Figure 2A**) and GDP (**Figure 2B**) bound simulations. A critical difference between the two nucleotide states is that GTP-bound simulations sample around one energy well, while the GDP- simulations sample between two energy wells. These two wells indicate conformations with Switch 1 folded into the protein (positive direction, predominantly occupied by WT) or folded out where it will no longer stabilize the active conformation. This result is concordant with the inactive nature of GDP-bound KRAS but is variably sampled by both hotspot and non-hotspot mutations. We quantified the time each mutation spent within these two energy wells (**Figure 2C**) and used the occupancy ratio to scale the mutations. For example, we found that WT, K5N, G12V, G13I, L23R, D30E, Q61K, and K117N exhibit low Switch 1 dynamics that favor active RAS conformations. Alternatively, variants such as G12D, Q22R, H27Y, F28L, P34Q, Q61R, and Y71D sample an extended Switch 1, favoring inactive RAS conformations. Thus, we find utility in the PC metric as a variant that overly favors the flexibility or rigidity of Switch 1 to be conducive to dysregulating RAS pathway signaling.

**Figure 2.**
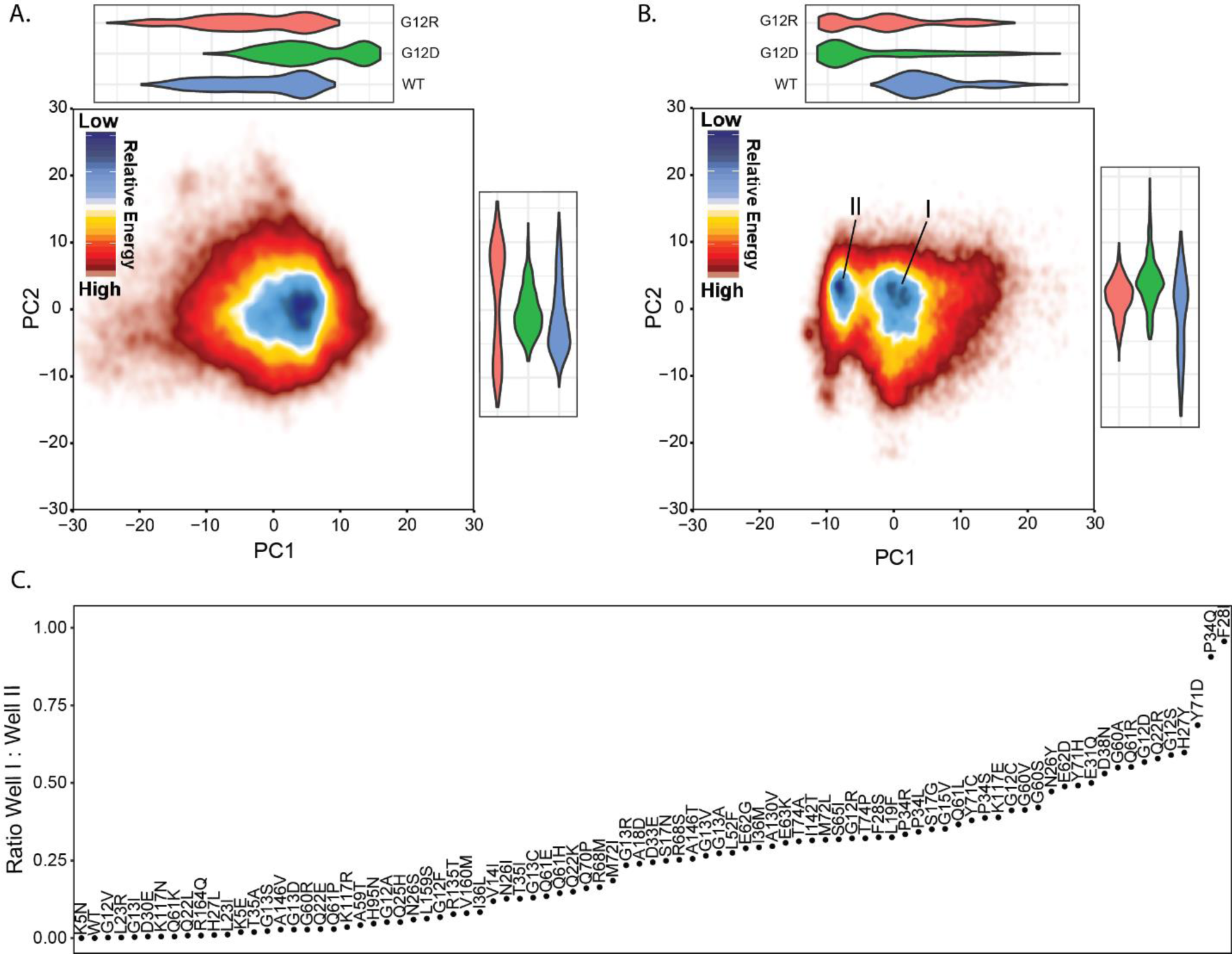
Principal Component Analysis reveals a duality in Switch 1 conformational dynamics. A) Free energy landscape of PC1 and PC2 of C_α_ coordinates within GTP simulations. The landscape is colored such that the dark blue region represents a higher density of coordinates, and the darker red represents a lower density of coordinates. Above and to the right of the landspace are violin plots representing the distributions of coordinate space for PC1 and PC2, respectively. The violin plots are colored such that WT is blue, G12D is green and G12R is red. B) Free energy landscape of PC1 and PC2 of C_α_ coordinates within GDP simulations. The landscape is colored such that the dark blue region represents a higher density of coordinates, and the darker red represents a lower density of coordinates. There are two different regions of blue level density where the right well is labeled energy well I and the left is marked energy well II. Above and to the right of the landscape are violin plots representing the distributions of coordinate space for PC1 and PC2 respectively. The violin plots are colored such that WT is blue, G12D is green and G12R is red. C) Ratio between GDP wells I and II labeled by variant and sorted according to the ratio. A ratio of 0 indicates a variant does not sample well II, and a ratio of 1 indicates a variant does not sample well I.

The utility of PCs in distinguishing between specific variants is further demonstrated by comparing G12D, G12R, and WT between GTP (**Figure 2A**) and GDP states (**Figure 2B**). We highlight these specific mutations because G12D is the most studied mutation, while G12R is gaining attention as a commonly observed in gastric cancers. The WT simulations exclusively sample stable closed Switch 1 conformations; G12D simulations exclusively sample extended Switch 1 out conformations, while G12R balances between them. Following this phenomenon is the Switch 1 RMSF of these three variants, which differs between these variants on a sequence basis (**Figure S1C**). G12R has high fluctuation values for amino acid positions 34 to 40, WT has high fluctuation values from 31 to 40, and G12D features high fluctuation values for amino acid positions 27 to 40. These specific details for how the phosphate-binding loop (p-loop) and Switches have altered dynamics are likely the underlying features for differences in intrinsic activities.

In summary, mutations with high Switch 1 flexibility are less WT-like and more G12D-like. This result is important because the flexibility of Switch 1 is indicative of a decrease in interaction between RAS and the nucleotide, a phenomenon that previous literature (28) describes as having the potential to cause nucleotide expulsion and or NF1 binding. As such, there is potential to estimate how much each variant favors the NF1 binding conformation by observing how similar Switch 1 dynamics are to the G12D simulations (**Figure 4C**). Therefore, in the following sections, we quantify ligand interactions and their associations with GAP and GEF binding conformations across mutations.

### Switch Conformational Dynamic Differences Group KRAS Mutations by Effector Binding Compatibility

We next compared the total difference between our MD ensembles and different solved KRAS structures in complex with physiologic binding partners. Because the Switch conformations regulate binding to GAPs and GEFs, we calculated their structural match to the GAP, NF1 (**Figure 3A**) and the GEF, SOS1 (**Figure 3B**). To compare these reference conformations, we calculated the RMSD between evenly selected conformations from our simulations. Our analyses identified consistent trends with mutations remaining nearby WT or favoring one of the binding competent conformations. Furthermore, the variants that favor the NF1 competent binding conformation over WT generally disfavor the SOS1 competent conformation and vice versa. The most extreme example is A18D which both favors the SOS1 binding conformation and disfavors the NF1 binding conformations greater than all other variants. By observing these trends across the variants, we separated the variants into groups based on deviation from WT. There are neutral variants which do not favor or disfavor any of the three conformations, compared to WT, comprised by 14/86 variants when comparing to NF1 (G12V, G13D, G13V, G15V, Q25H, N26Y, D33E, T35A, T35I, A59T, Q70P, R135T, A156V) and 14/86 variants for when comparing to SOS1 (G13R, G13I, V14I, G15V, Q22L, L23I, T35I, N26S, A59T, G63M, Y71C, H95N, A146V, L159S). The other variants, which are more than 1σ from WT, favor or disfavor one or more reference conformations depending on which nucleotide is bound, which offers insight into how these variants cause disease. For example, K5N has a similar deviation to WT from the SOS1 binding conformation when GDP is bound but is 3σ from WT when GTP is bound, indicating that K5N lacks affinity for the SOS1 binding conformation only when GTP is bound. Mutant proteins in these alternative groups have a higher propensity to maintain specific binding competent conformations instead of changing between conformations, predicting the mechanism by which these variants dysregulate the RAS pathway to cause disease.

**Figure 3.**
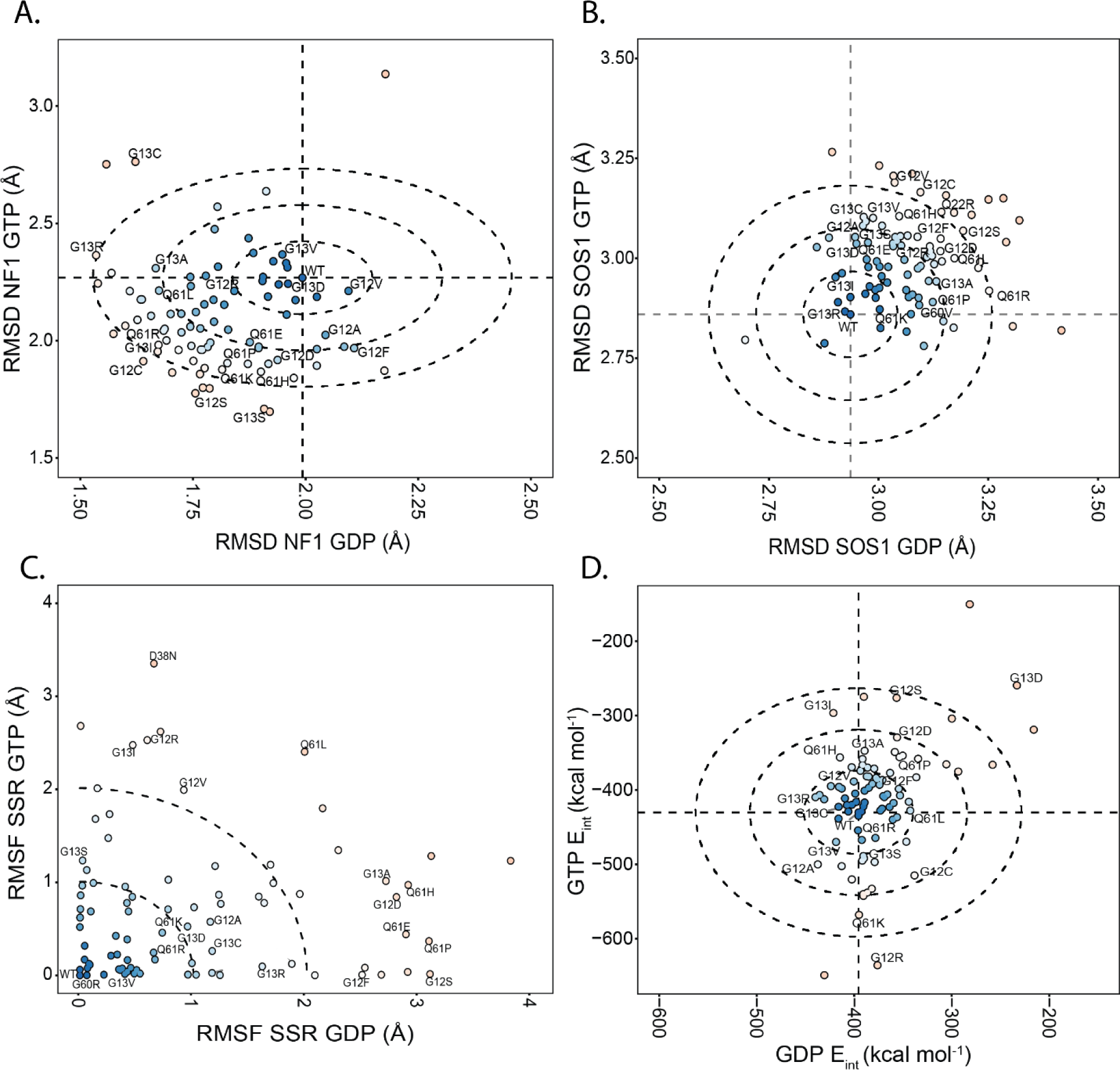
KRAS mutations exhibit significant differences in effector binding conformational dynamics. A) The replicate-averaged mobility of each amino acid displays the largest flexibility for the two Switches, yet with high mobility that is variable across mutations, in the allosteric lobe. Dotted lines mark the WT values. B) The replicate-average difference from solved NF1 binding competent KRAS pose (PDB: 6OB2). C) Sum of Square Residuals (SSR) from the average RMSF between all variants on a per amino acids basis. D) Variants feature differing stabilities concerning the presence of the nucleotide in the KRAS binding pocket. The horizontal and vertical lines represent the average nucleotide interaction energy value for WT. The concentric circles represent how many standard deviations away variants are from the WT. The inner circle represents one standard deviation, the middle circle represents two standard deviations, and the outer circle represents three standard deviations. Each variant is also colored according to its distance from the WT. Variants colored in blue feature similar E_int_ values to the WT. Red color variants represent variants that are far from the WT.

RMSD is an excellent scoring method for assessing overall conformational change but does not provide a complete picture regarding overall trends in conformational dynamics. Therefore, we first quantified per-mutation differences in Switch and allosteric region flexibilities (**Figure S2**). Globally, Switch 1 and 2 have the most flexibility in our system, concordant with the conventional understanding of KRAS dynamics and data from across all available experimental structures (**Figure 1C** and **1D**). We used the different degrees of flexibility among mutations to subdivide mutations by their flexibility patterns, within and between Switches, and across the C-terminal allosteric lobe. Differences in dynamic patterns were calculated using the SSR from the WT profile (**Figure 3C**). This approach identified variants with similar dynamics to WT and variants that deviate. In GTP simulations (**Figure 3C**), 42/86 have |SSR^MT^ - SSR^WT^| < 0.5, indicating similar flexibility profiles. Likewise, in GDP simulations (**Figure 3C**), 41/86 are within 0.5 of SSR^WT^. There is an overlap of 21/86 variants (24%) within 0.5 SSR in both the GDP and GTP bound states and 45 (52%) within 0.5 in either. Reciprocally, 41/86 (48%) variants demonstrate increased variability in conformational dynamics compared to WT. Also, mutants with high sum of square residuals (SSR) in both GDP and GTP simulations, such as N26I, Q61L, and A146T, overly favor the SOS1 binding conformation regardless of the nucleotide-bound state. SSR values and mutational groups are listed in **Tables S2**. We anticipate these mutations alter cellular signaling primarily through changes to their intrinsic dynamics, which dysregulates coupling to effector proteins and thereby defines their role in human diseases.

### Interaction Energy Elucidates Differing Nucleoside Binding Responses on a Multi-Variant Basis

For KRAS, the interactions between the nucleotide and the KRAS switch regions are well-accepted as important for proper protein function. One crucial use of MD is calculating the specific properties concerning the non-bonded interactions throughout the protein. As such, we leveraged this critical aspect of MD and calculated the interaction energy between the nucleotide and KRAS protein (**Figure 3D**). The distribution of values on a per-variant basis divided the variants into distinct groups based on the difference from the WT. Most variants (41) are within 1σ of WT, which indicates little change in ligand interaction energy. The variants between 3σ and 4σ from WT are T3A, G12S, Q13I, A18D, Q25H, Q61K, Q70P, and K117E. The variants ≥4σ from WT include G12R, G13D, G15V, Q60R, and K117N. Thus, KRAS mutations fall into a stabilizing, neutral, and destabilizing spectrum. For example, E31Q is 1.5σ from the WT interaction energy for both GTP and GDP, indicating the protein has a more energetically favorable interaction with both nucleotides. Alternatively, Q25H has a similar GDP interaction energy to WT and a less stable GTP interaction energy at almost 3σ from WT. These results indicate that Q25H has favorable interactions with GDP but less favorable interactions with GTP, which likely functionally complement each other to lead to reduce intrinsic GTPase activity. The energetic scores for all proteins are available in **Table S2**. In summary, relative nucleotide stability informs which variants have overly unfavorable or favorable magnitudes of nucleotide interactions, which have a strong potential to predict how variants can mishandle the nucleotide and dysregulate the RAS signaling pathway.

### Biophysical Interpretation of Molecular Dynamics Scores Describes Mutational Functions

Above, we assessed individual MD-based metrics that identify differences in conformational sampling across KRAS mutant proteins and predict features of their intrinsic differences. Now, we biophysically annotate the PC scores and interpret our simulations in the context of precise observations established from the extensive experimental literature on RAS. First, we identified a set of ten pairwise distances within KRAS that are either markers for biochemical activities or critical structural features of distinct RAS conformations according to biophysical measurements including from NMR spectroscopy (**Figure 4B**) (3,27,28,37). These ten distance pairs fall into four themes associated with different hotspot mutations (**Table 1**). Precisely, they quantify the extent of opening Switch 1 and Switch 2 from the p-loop, the coordination of movements between Switches, and the electrostatic coordination of the magnesium cation accompanying GTP. MD-based PCs capture switch motions, the basis for the four themes (**Figure 4B**) and strongly correlate with the ten distance monitors (**Figure 4C**). For example, the distance monitor Y32 to Y59, which describes the interconnectedness between Switch 1 and Switch 2, correlates -0.75 with both PC1 and PC2, demonstrating that when variants favor the negative directions of PC1 and PC2, the Switch regions separate. Additionally, there is nearly a reciprocal relationship between PC1 in the GDP and GTP states, where Switch 1 has the opposite movement direction as the enzyme shifts between active and inactive conformations. In the GTP simulations PC1 is anticorrelated -0.8 with distance monitors G12 to T35 and Q61 to T35. This trend is reversed in the GTP simulations demonstrating fundamental differences in how the nucleotide affects KRAS dynamics. These data confirm the functional and biophysical nature of our simulations and establish their biophysical interpretation.

**Table 1:** Description of previously characterized structural hallmarks of KRAS activity.

| Amino Acid Pair Distance | Description | Mutation Studied | CC<sub>GTP</sub> | CC<sub>GDP</sub> |
| --- | --- | --- | --- | --- |
| Q61 to D92 (28,37) | Hbond between $\alpha 2$ helix and $\alpha 3$. Lost in mutation, favoring loop-out | G12D, WT | PC1 + .1<br>PC2 + .8<br>PC3 + .1 | PC1 + .1<br>PC2 + .5<br>PC3 - .4 |
| E62 to H95 (28) | Hbond between $\alpha 2$ helix and $\alpha 3$. Lost in mutation, favoring loop-out | G12D, WT | PC1 + .0<br>PC2 + .7<br>PC3 + .1 | PC1 + .0<br>PC2 + .5<br>PC3 - .6 |
| A11 to Q61 (28) | Hbond between p-loop and Switch 2 and $\alpha 2$. Lost in mutation, favoring loop-out | G12D, WT | PC1 + 0<br>PC2 + .6<br>PC3 + .5 | PC1 + .1<br>PC2 + .8<br>PC3 - .2 |
| G12 to Q61 (28) | Hbond between p-loop and switch 2 and $\alpha 2$. Lost in mutation, favoring loop-out | G12D, WT | PC1 - .2<br>PC2 + .3<br>PC3 + .7 | PC1 + .0<br>PC2 + .8<br>PC3 + .1 |
| I36 to A59 (3) | Hbond between $\beta 2$ and $\beta 3$ ladder which are between the switches. Mutation destabilizes Switch 1 and Switch 2 coordination | G12D, G13D, WT | PC1 - .8<br>PC2 - .3<br>PC3 + .1 | PC1 + .8<br>PC2 + .1<br>PC3 + .1 |
| Y32 to Y40 (3) | Hbond within switch as well as ring stacking. G13D favors intermediates not used by WT | G12D, G13D, WT | PC1 + .1<br>PC2 + .1<br>PC3 + .1 | PC1 - .4<br>PC2 - .1<br>PC3 + .0 |
| Y32 to A59 (3,37) | Describes the interconnection between switch 1 and switch 2. G13D favors A59 replacing Mg<sup>2+</sup>, mimicking HRAS:SOSs | G12D, G13D, WT | PC1 - .7<br>PC2 - .6<br>PC3 + .1 | PC1 + .3<br>PC2 + .1<br>PC3 + .0 |
| G12 to T35 (27) | T35 in Hbond network among nucleotide and Q61. Network destabilized by mutation | G12V, Q61L | PC1 - .7<br>PC2 + .1<br>PC3 + .1 | PC1 + .9<br>PC2 + .2<br>PC3 + .1 |
| Q61 to T35 (27) | T35 in Hbond network among nucleotide and Q61. Network destabilized by mutation | G12V, Q61L | PC1 - .7<br>PC2 - .5<br>PC3 + .2 | PC1 + .8<br>PC2 + .1<br>PC3 + .3 |
| S17 to T35 (27,37) | Failure to bind Mg<sup>2+</sup> and coordinate T35 | G12V, Q61L | PC1 + .2<br>PC2 + .2<br>PC3 + .1 | PC1 + .3<br>PC2 + .1<br>PC3 + .0 |

**Figure 4.**
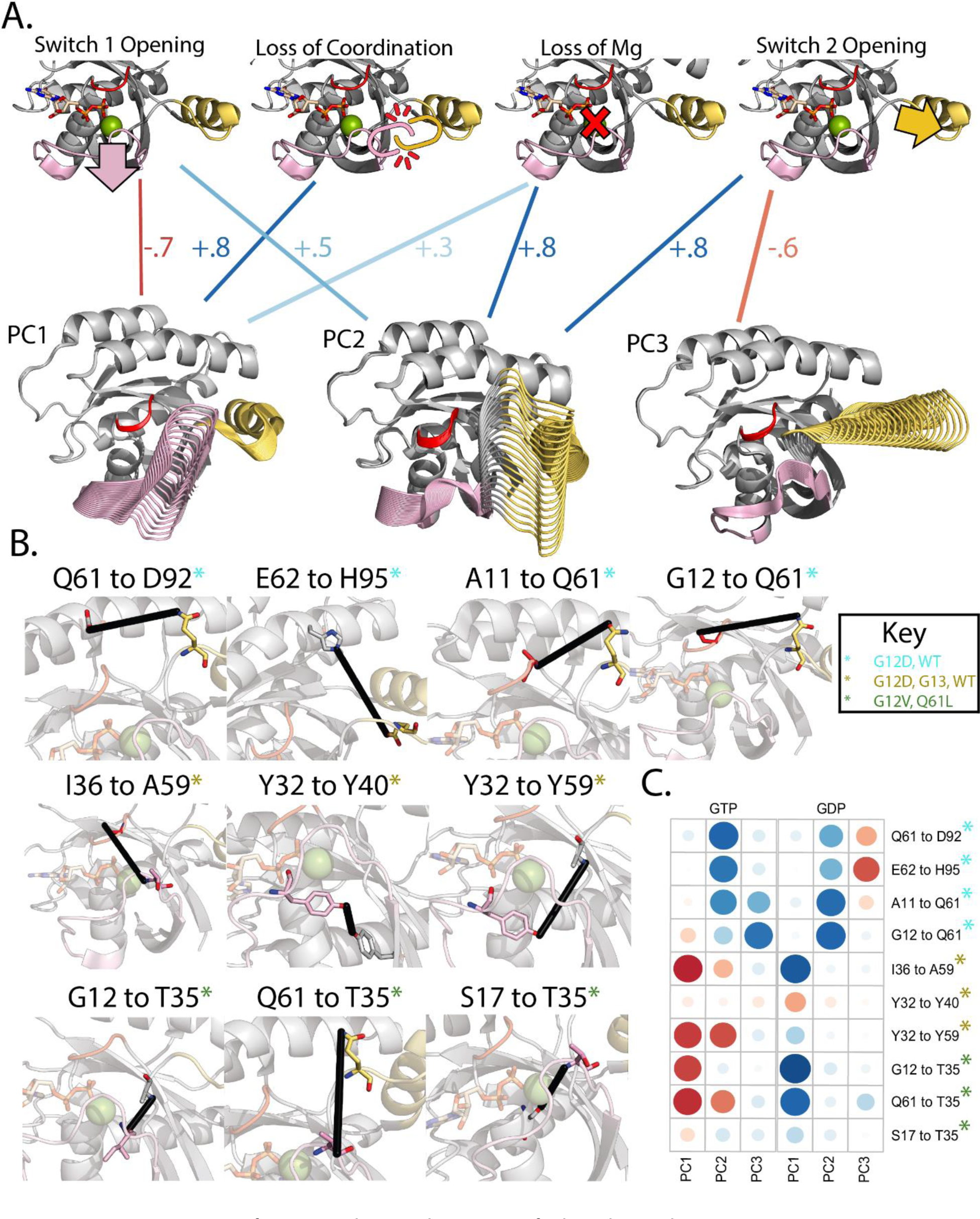
PC movements from simulations have specific biophysical interpretations. A) Leveraging the deep history of RAS study, we identified ten distance measurements that have been experimentally characterized and used as monitors to detect four difference biophysical phenomena (Table 2). Taken together, the first three PCs strongly associate with four biophysical phenomena. B) Display of each distance monitor used to describe the conformational change. Amino acids in each monitor are rendered in sticks while the remainder of the KRAS structure is rendered in cartoon. To display a potential interaction between these monitors a black line is drawn between the R-groups of relevant amino acids. C) Full correlation plot between the PCs and the distance monitors. Red circles represent negative correlations while blue circles represent positive correlations. The size represents the magnitude of correlation strength where a larger circle is a stronger correlation.

To determine relationships among all MD-based metrics (spanning RMSF, PCs, interaction energies, and distance monitors) we computed a complete correlation matrix (**Figure S2A** and **Figure S2B**), revealing relationships among these metrics. As previously discussed, PC scores correlate with groups of distance monitors. Furthermore, the PCs and these distance monitors correlate with groups of RMSD and RMSF metrics depending on the binding competent conformations and effector regions the PCs correspond to. This correlation structure is subdivided by metrics that strongly affect Switch 1 and metrics that affect Switch 2. Metrics correlating with Switch 1 include: global RMSF, RMSF of Switch 1, RMSD NF1, RMSD SOS1, RMSD DESRES, G12 to I35, I36 to A59, Y43 to Y59, and G61 to T35. Switch 2 includes: RMSF of Switch 2, RMSD NF1, RMSD DESRES, A11 to Q61, G12 to Q61, Q61 to D92 and E62 to H95. We also generated a heat map of the MD metrics on a per-variant basis to gain insight into which variants have similar score profiles (**Figure S2C**). K-means clusters of the heatmap data divide the mutations with similar SSR profiles, with about half of the mutations demonstrating significant changes in multiple scores simultaneously. For example, a slight change in Switch 1 or 2 has the potential to cascade and cause larger conformational changes, which are observed in unison by our MD metrics. The correlation matrix provides a complete summary of the interconnectedness between our MD metrics and the multi-faceted effects of each mutation on KRAS intrinsic dynamics.

### Biophysical Measurements Confirm the Uniqueness of Each Mutated Protein

We developed a uniform high-throughput platform for recombinantly expressing and purifying human KRAS mutants from *E. coli* (see Methods). We successfully purified 65 of the 86 mutant proteins; the 21 not purified are K5E, G13I/V, V14I, G15V, S17G, A18D, L19F, Q22L, L23R, N26I, T35I, K117E/N/R, and A146V. We used thermostability measurements to determine relative similarities among the 65 KRAS mutations and their relative propensity for activating (GTP-stabilized) or inactivating (GDP-stabilized) conformations. After optimizing buffer conditions, WT KRAS measurements were highly reproducible with the GDP-bound form melting transition at 62.8 ± 0.9°C, and 55.6 ± 1.4°C when bound to the non-hydrolyzable the GTP-analog, GMP-PNP (referred to as “GTP state” herein). Thus, WT-KRAS is more stable in the inactive GDP-bound form by 7.2°C. Temperatures of unfolding were highly similar between the two ligands and across mutations (p = 26x10^-7^;), demonstrating a consistent pattern and system balance.

Thermal melting temperatures of KRAS mutations spanned a wide range of values from highly destabilizing (40°C) to stabilizing (65°C; **Figure 5A**). Differences between GDP and PNP were bookended on one end by mutations with minor differences such as Q61R (ΔΔT_m_^GDP-PNP^=0.3°C) and G12V (ΔΔT_m_^GDP-PNP^=1.4°C), and on the other end by mutations with stark differences such as I36L (ΔΔT_m_^GDP-PNP^=15.8°C) and G60V (ΔΔT_m_^GDP-PNP^=16.2°C). Hotspot mutations occur across this range, such as G12F (ΔΔT_m_^GDP-PNP^=14.0°C) and G12D (ΔΔT_m_^GDP-PNP^=12.4°C). Further, observing that some mutations stabilize the active conformation while others destabilize the inactive form, we calculated the overall activation stability change as ΔΔT_m_* = ΔT_m_^PNP^ - ΔT_m_^GDP^. KRAS mutations ranged in ΔΔT_m_* from inactivating, exemplified by Q61R (ΔΔT_m_* = 6.99°C), G12V (ΔΔT_m_* = 5.88°C), and Q22E (ΔΔT_m_* = 5.62°C), to activating, exemplified by G60V (ΔΔT_m_* = -8.95°C), I36L (ΔΔT_m_* = -8.59°C), and G60R (ΔΔT_m_* = -8.04°C). Q61R was expected to have the greatest difference in activated compared to inactivated KRAS stability, yet both nucleotide-bound forms were destabilized. Conversely, compared to G60V, with the greatest destabilization of the activated KRAS and little effect on the inactivated form. Mutations such as G12V stabilize both states but stabilize the GTP state much more than the GDP state. Other mutations such as G13C/R/S and G12C destabilize both states equally. At certain non-hotspot sites, such as G60, loss of the native amino acid has a larger effect on stability than differences in the alternate amino acids; G60A/R/V all have minor effects on GDP state (ΔΔT_m_ = 0.5 ± 0.7°C), but significantly destabilize the GTP state (ΔΔT_m_ = -5.8 ± 3.4°C). The most extreme destabilization occurs for A146T, but the change is greater for the GDP state (-16.7°C) than for the GTP state (-11.5°C). Across our thermostability measurements, the mutational landscape of KRAS produces proteins with a spectrum of nucleotide-dependent stabilities that we anticipate alter intrinsic activity.

**Figure 5:**
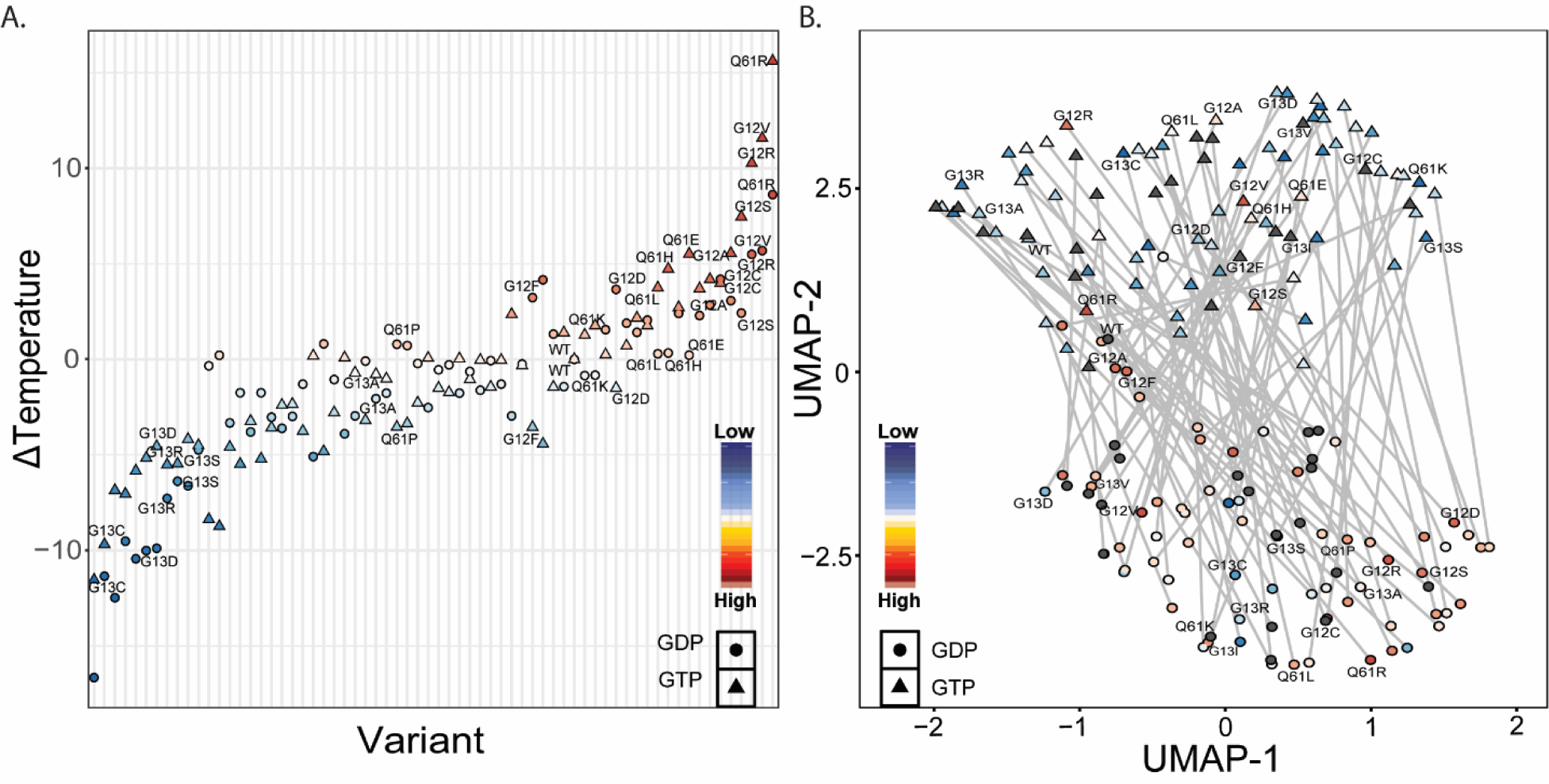
Using combined of metrics allows dimensionality reduction that mirrors T_m_ Data. A) Mean T_m_ difference from WT on a per variant basis sorted by difference from WT. Symbols are as follows; circles represent GDP-bound variants and triangles represent GTP-bound variants. Each symbol is colored according to their difference from WT, where blue symbols feature a lower value than WT and red symbols feature a higher value than WT. B) UMAP dimensionality reduction calculated using all MD metrics discussed in this study. Each variant is colored according to T_m_ as seen in A. Variants colored in grey have simulation data but do not have T_m_ data.

### Computed Features from Molecular Simulations Predict Biophysically Meaningful Values

Again, using our full complement of scores, UMAP embedding was used to compare the relative similarities across all variants (**Figure 5B**). The results are promising because the UMAP captures the distinction between GTP and GDP states, with GDP states occupying one cluster, GTP states occupying a second cluster, and little overlap between clusters. We projected our melting temperature measurements onto the UMAP space, revealing that the GDP variants have a higher melting temperature compared to the GTP variants, yet each following a gradient; within GTP-bound simulations, we observe a linear spectrum with the most stabilized mutations on one side and least stable on the other. These data affirm that, even though melting temperature is a single composite measurement, our collection of MD-based scores captures features relevant to explain experimental results. Additionally, we took the PC standard scores and colored the UMAPs by the values of those scores. Interestingly, we found a split in the PC1 standard scores that closely resembles the split in melting temperature data (**Figure S3**). Conversely, the PC2 and PC3 standard scores appear unrelated to the melting temperature values. As such, we conclude that switch 2 confirmation has a lower effect on overall KRAS conformational stability when compared to Switch 1 conformation.

### Grouping of Variants Elucidates the Utility of MD Scores in Categorizing Variants by Their Conformational Dynamics

So far, we have described several dynamics-based scores that follow established KRAS biophysical metrics and have utility in identifying critical differences between variants. Using diverse metrics simultaneously, we can more robustly subdivide and characterize groups of KRAS mutations. To better accomplish this, we defined six groups using all calculations (**Figure S2**) to generate meta-classifications, which we define as clusters of KRAS mutations that share characteristics across all scores. Individual scores were used to classify mutations as WT-like (within 1σ), less than WT (< -1σ), and greater than WT (> 1σ), making 90 total meta-classifications (**Figure 6**). Every group has at least one hotspot variant except for GTP Group 5, the smallest group with five variants. GDP Groups 3 and 4, as well as GTP group 6, all have one hotspot variant. GDP Group 6 and GTP Group 3 have the most hotspot variants with six and seven, respectively. All the other groups have three or four hotspot variants (**Table 2**). The spread of hotspot mutations across these profile groups indicates the importance of characterizing non-hotspot genetic variants and their role in congenital diseases and human cancers.

**Figure 6:**
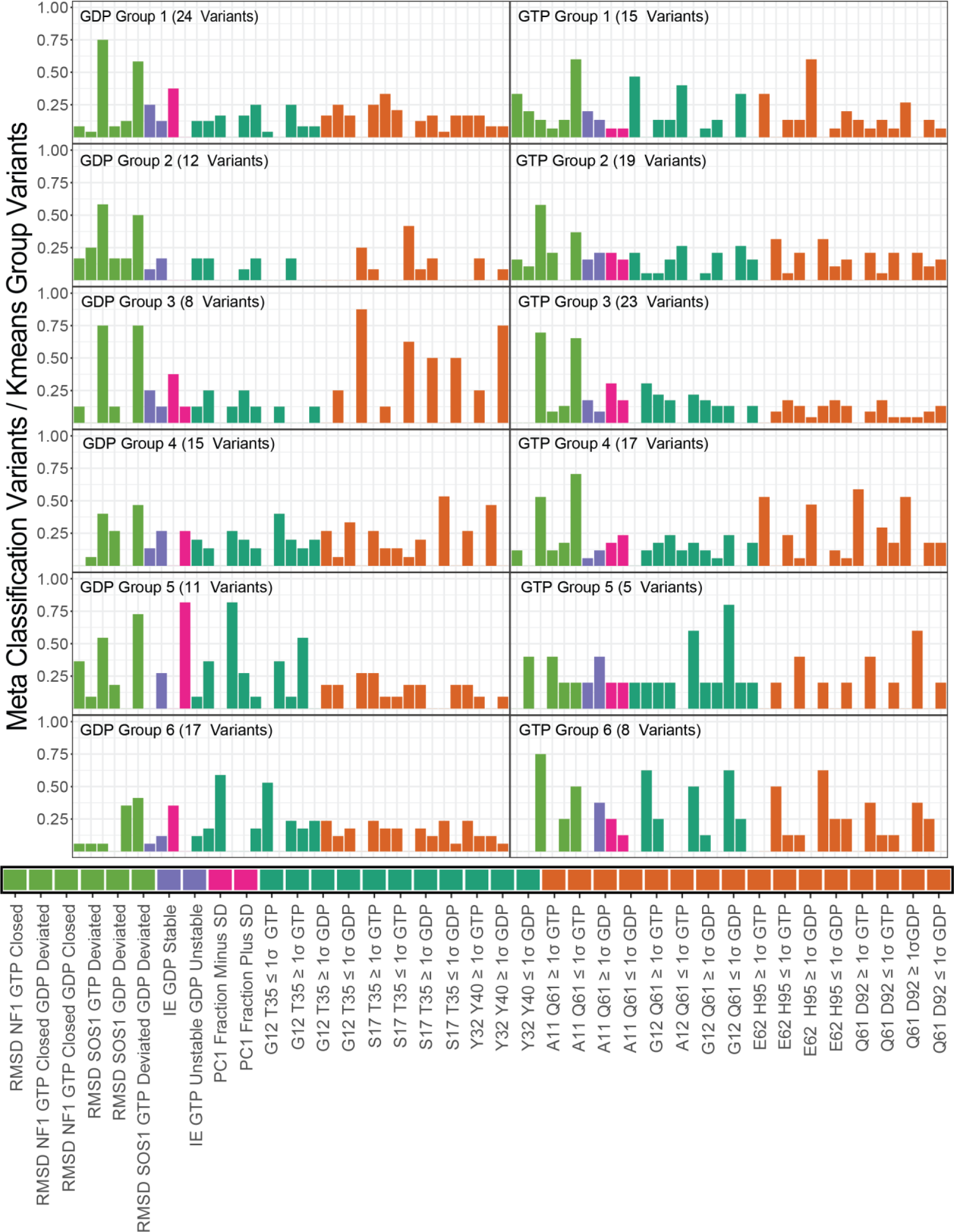
Rankings of Different Meta Classifications by Occupancy Across Clustering Groups. We define a Meta-class as the K-means groups across the 90 metric-specific classifications, which were then filtered down to 37 classifications where each classification is in at least one of the top 10 classifications for each K-means Group. The bars are ordered by the type of score that each Meta Classification comes from. The score type is indicated by the lengthwise bars below each plot. Green indicates RMSD, Purple indicates Interaction Energy, Magenta Indicates PC1 fraction, Teal indicates Switch 1 Distance Monitor, and Orange indicates a Switch 2 Distance Monitor. The Distance monitor Meta-classes that either.

**Table 2:** Variant Identity of each Kmeans Group

| Kmeans Group | Identity |
| --- | --- |
| GDP Group 1 (24 variants) | K5E, **G12R, G13S**, Q22K, Q22L, L23I, N26I, N26S, P34S, T35A, I36L, G60R, **Q61H**, Q61P, E62G, E63K, S65I, R68M, M72I, T74A, H95N, K117N, A146V, R164Q |
| GDP Group 2 (12 variants) | **G12C, G13A, G13C, G13R**, V14I, F28S, P34L, P34R, G60S, R68S, K117E, K117R |
| GDP Group 3 (8 variants) | **G13I**, L19F, Q22E, H27L, Q61K, E62D, Q70P, T74P |
| GDP Group 4 (15 variants) | **G12D**, G15V, S17N, A18D, N26Y, E31Q, T35I, L52F, G60V, Y71C, Y71H, M72L, A130V, I142T, A146T |
| GDP Group 5 (11 variants) | **G12S**, S17G, Q22R, H27Y, F28L, P34Q, D38N, G60A, **Q61L, Q61R**, Y71D |
| GDP Group 6 (17 variants) | **WT**, K5N, **G12A, G12F, G12V, G13D, G13V**, L23R, Q25H, D30E, D33E, I36M, A59T, **Q61E**, R135T, L159S, V160M |
| GTP Group 1 (15 variants) | **G12R, G13C, G13D**, N26I, D33E, T35I, D38N, **Q61L**, E63K, Q70P, M72L, K117E, A130V, A146T, V160M |
| GTP Group 2 (19 variants) | **WT**, K5E, **G13A, G13R**, V14I, G15V, S17G, S17N, Q22E, Q22L, H27L, H27Y, P34Q, I36L, L52F, A59T, **Q61R**, E62G, R68M |
| GTP Group 3 (23 variants) | **G12C, G12D, G12F**, G13I, G13S, Q22R, L23I, L23R, F28S, D30E, E31Q, P34R, I36M, **Q61E, Q61H, Q61K, Q61P**, E62D, T74A, K117R, I142T, L159S, R164Q |
| GTP Group 4 (17 variants) | **G12A, G12V, G13V**, L19F, Q22K, N26Y, P34S, T35A, G60A, G60R, G60S, G60V, S65I, Y71D, Y71H, M72I, R135T |
| GTP Group 5 (5 variants) | A18D, Q25H, F28L, P34L, A146V |
| GTP Group 6 (8 variants) | K5N, **G12S**, N26S, R68S, Y71C, T74P, H95N, K117N |

Each profile has unique features. Indeed, the different groups can readily be distinguished by their meta classifications. For instance, GTP Group 4 has a higher overlap with distance monitors than GTP Group 2, even though the two groups have a similar number of variants. Investigating the top 5 meta classifications for each profile described the types of dynamics seen within each variant group (**Table 3**). For example, observing the patterns of meta-classes within GTP Group 4 we discern several points of crucial information. Primarily, the RMSD classification indicates that the variants are conformationally close to the NF1 binding competent conformation and far from the SOS1 binding competent conformation. Three distance monitors explain the precise mechanism underlying this conformational restriction that favors the NF1 binding conformation. E62-H95 and Q61-D92 are monitors for Switch 2, and both indicate greater flexibility in this group of variants. The A11-Q61 monitor describes Switch 2 movement away from the nucleotide. This is because A11 is within the p-loop involved in nucleotide interactions, and Q61 is within Switch 2. As such, we can summarize GTP Group 4 as a group of variants that favors NF1 binding and flexibility of Switch 2. Using our meta-classification information (**Table 3** and **Figure 6**), similar descriptions are evident for every group. Therefore, we display the capacity of molecular dynamic data to provide a systematic interpretation of genetic variations, enabling a functional understanding of the KRAS mutational landscape.

**Table 3:** Summary of Top 5 Input Features for Distinguishing each Meta Classification

| Cluster Group | Meta Class Rank |  |  |  |  |
| --- | --- | --- | --- | --- | --- |
|  | 1 | 2 | 3 | 4 | 5 |
| GDP Group 1 (24 variants) | RMSD NF1<br>GTP Closed<br>GDP Closed | RMSD SOS1<br>GTP Deviated<br>GDP Deviated | PC1 Fraction<br>Minus SD | G12 Q61<br>Below GTP | G12 Q61<br>Above GTP |
| GDP Group 2 (12 variants) | RMSD NF1<br>GTP Closed<br>GDP Closed | RMSD SOS1<br>GTP Deviated<br>GDP Deviated | G12 Q61 Below<br>GDP | A11 Q61<br>Below GDP | RMSD NF1<br>GTP Closed<br>GDP Deviated |
| GDP Group 3 (8 variants) | A11 Q61<br>Below GDP | Q61 D92<br>Below GDP | RMSD SOS1<br>GTP Deviated<br>GDP Deviated | RMSD NF1<br>GTP Closed<br>GDP Closed | G12 Q61<br>Below GDP |
| GDP Group 4 (15 variants) | E62 H95<br>Above GDP | Q61 D92<br>Above GDP | RMSD SOS1<br>GTP Deviated<br>GDP Deviated | S17 T35<br>Below GDP | RMSD NF1<br>GTP Closed<br>GDP Closed |
| GDP Group 5 (11 variants) | G12 T35<br>Below GDP | PC1 Fraction<br>Plus SD | RMSD SOS1<br>GTP Deviated<br>GDP Deviated | Y32 Y40<br>Above GDP | RMSD NF1<br>GTP Closed<br>GDP Closed |
| GDP Group 6 (17 variants) | G12 T35<br>Above GDP | S17 T35<br>Above GDP | RMSD SOS1<br>GTP Deviated<br>GDP Deviated | PC1 Fraction<br>Minus SD | RMSD SOS1<br>GDP Deviated |
| GTP Group 1 (15 variants) | G12 Q61<br>Above GTP | RMSD SOS1<br>GTP Deviated<br>GDP Deviated | G12 T35 Above<br>GTP | S17 T35<br>Above GTP | A11 Q61<br>Above GTP |
| GTP Group<br>2 (19<br>variants) | RMSD NF1<br>GTP Closed<br>GDP Closed | RMSD SOS1<br>GTP Deviated<br>GDP<br>Deviated | G12 Q61 Below<br>GTP | A11 Q61<br>Below GTP | Y32 Y40<br>Above GDP |
| GTP Group<br>3 (23<br>variants) | RMSD NF1<br>GTP Closed<br>GDP Closed | RMSD SOS1<br>GTP Deviated<br>GDP<br>Deviated | G12 T35 Below<br>GTP | PC1 Fraction<br>Minus SD | S17 T35<br>Below GTP |
| GTP Group<br>4 (17<br>variants) | RMSD SOS1<br>GTP Deviated<br>GDP Deviated | E62 H95<br>Above GTP | Q61 d92<br>Above GTP | A11 Q61<br>Above GTP | RMSD NF1<br>GTP Closed<br>GDP Closed |
| GTP Group<br>5 (5<br>variants) | Y32 Y40<br>Above GTP | Q61 D92<br>Below GTP | S17 T35 Below<br>GTP | E62 H95<br>Below GTP | A11 Q61<br>Below GDP |
| GTP Group<br>6 (8<br>variants) | RMSD NF1<br>GTP Closed<br>GDP Closed | G12 Q61<br>Below GTP | Y32 Y40 Above<br>GTP | G12 T35<br>Below GTP | A11 Q61<br>Below GTP |

### Patterns From Dynamics Calculations Across Mutations Match Existing Experimental Data

We consulted our established RAS resource (3,27,28) for groups of mutations that share experimentally derived features and compared them to our results described above. Two major hotspots, G12D and G13D, have been studied extensively via biochemical, biophysical, and cellular experiments. These previous studies revealed that both favor non-WT-like conformations with G12D favoring the GTP-bound, Raf-bound state. In contrast, G13D favors this state to a lesser degree due to its destabilization of the binding pocket and the development of additional intermediates. G12D and G13D lie on nearly opposite sides of our UMAP space (for GTP and GDP forms), supporting their divergent effects on structure and dynamics in simulation. Additionally, the G12V and G13V classic hotspot mutations lie close in UMAP space to G12V in the active GTP-bound state yet diverge in the GDP-bound state. Heterogeneity in fibroblast and epithelial cell response to distinct Q61 mutations (38) match our results in that Q61E/P display a different pattern of dynamics-based scores compared to Q61H/L/R, the latter of which more closely resemble G12D. Yet, in our analysis and previous experiments, Q61R remains closer to Q61P in character than Q61H/L. Non-hotspot mutations are less studied experimentally; in our assessment, non-hotspot mutations exhibited an even more diverse array of structure- and dynamics-based changes than hotspot mutations.

## DISCUSSION

We have uniformly characterized a large and diverse repertoire of 86 KRAS mutations, spanning the alterations observed in congenital diseases and human tumors. Our simulation-based approach to representing the effects of genomic mutations on encoded 3D protein structures and their dynamics is scalable to hundreds of mutations, sensitive to details of the molecular environment (e.g. GDP versus GTP), and captures patterns identified through experimentation. This level of mechanistic specificity and molecular detail is currently lacking in genomics. Thus, our approach extends our previous work and achieves increased resolution for genomic data interpretation.

In previous work, we advanced the field of translational genomics by using calculations of 3D translated gene products, rather than the gene sequence, for better interpreting cancer-causing gene mutations and inter-individual genetic variation (4). We also used experimental structures across the RAS GTPase family to scale the interpretation of human genetic variation by leveraging structure bioinformatics into a thousand variants (5). In the current study, we explored the utility of 3D dynamic information for extending structural bioinformatics for genomics. We developed a series of dynamics scores that capture established experimental biophysical markers and used them to define groups of variants with related effects. Our approach of standardizing PC-based scores to WT dynamics was particularly helpful in matching with experiments. The strong correlation between the PC standard scores and biophysical markers allows for an interpretation of how different types of conformational change will cause different effects on the intrinsic activity of this key enzyme and signaling regulator. High-level trends across the structure and dynamics-based calculations also correlated with melting temperature values. As such, we have increased confidence in the utility of dynamics scores in differentiating between genomic variations. Indeed, these results establish the potential to use dynamics scores on a genomic basis to distinguish disease variants and their potentially divergent mechanisms, in future studies.

While the field now acknowledges heterogeneity in enzymatic and cellular outcomes across KRAS mutations, the ability for computational approaches to elucidate details of intrinsic dynamics may not be universally accepted. As data science tools continue to advance, their role in hypothesis generation, testing, and functional inferences similarly advances. Given the increasing accessibility of high-performance computing, ingenuity in algorithms, and complexity of analytic resources, studies such as the current work will establish a new level of mechanistic information about mutant proteins. This data is applicable to all human body tissues and cellular contexts, which can be applied to systems modeling to increase predictive and inferential power.

With any simulation-based study naturally comes the question of adequate sampling. The computational power used for simulations must be realistically balanced with the scope of molecular phenomena expected, for such methods to scale on a genomic basis. In this study we performed medium-length simulations to assess a baseline for the comprehensive study of inter-individual KRAS variation, and as a pilot for their application on a larger basis. Indeed, we were able to observe important dynamics features from the current extent of sampling. Yet, we expect our future studies to focus on enhanced sampling techniques over longer classic MD simulations to better enumerate the mutation-specific conformations and dynamics, which will likely elucidate additional details and nuance for the specific intrinsic molecular dysregulation. Likewise, the current simulations carry clear biophysical information yet do not precisely recapitulate changes in T_m._ Enhanced sampling will better elucidate the ensemble properties that reflect thermostability. While thermostability is important, especially for KRAS and as it applies to stabilizing or destabilizing the active and inactive states, thermostability may provide limited information regarding the precise dimensions of enzyme function. Protein dynamics are too complex to be adequately described by a single value in an experiment or MD. The strong correlations between our analysis and established experiments indicate merit in medium-length simulations for genomics, and patterns of KRAS dysregulation that span hotspot and non-hotspot mutations.

In summary, we successfully developed a new scalable workflow capable of functionally grouping genomic variants using MD scores. In using established experimental findings, we contextualized and validated our data. MD has been used mainly for studies that are biophysical in nature, but our workflow presents an expanded scale of MD, utilizing the method in the context of MD utilized in genomics. Our workflow presents an expanded scale in the context of MD utilized in genomics. Furthermore, MD is uniquely advantageous in its utility as a method due to its capacity to score variants in a uniform manner. As such, we will continue improving our workflow to increase the ease and commonality of using MD to explain inter-individual genetic variation, which may one day be applied to not only explain disease mechanisms but also to inform treatment with mutation-specific therapeutics targeting mutationally restricted conformations.

## DATA AVAILABILITY

All calculations generated in this study are included in the supplemental data files. Additional raw data can be provided by the authors upon reasonable request.

## AUTHOR CONTRIBUTIONS

RU and MZ Conceptualized the study. BR, DJ, AM, BV, and MZ contributed methodologies, resources, and software. BR and MZ wrote the original draft. BR, NH, JW, GE, and EL performed formal analysis and investigations. DJ, AM, GL, BS, BV, RU, and MZ provided supervision and administration. GL, BS, BV, RU, and MZ acquired funding for the project. All authors contributed to critical review, editing, and approved the final draft.

## FUNDING

This work was supported in part by the Linda T. and John A. Mellowes Center for Genomic Sciences and Precision Medicine at the Medical College of Wisconsin (<u>RRID:SCR_022926</u>). Supported by a grant from the State of Wisconsin Tax Check-Off Program for Cancer Research and the Medical College of Wisconsin Cancer Center. This research was completed in part with computational resources and technical support provided by the Research Computing Center at the Medical College of Wisconsin. This project is funded in part by the Advancing a Healthier Wisconsin Endowment at the Medical College of Wisconsin. National Institutes of Health Grants R01 DK52913, R01 CA247898, and R35 GM128840 supported this work.

## CONFLICT OF INTEREST

BV declares ownership interests in Protein Foundry and XLock Biosciences.

