## Supplementary material for "Beyond Structural Bioinformatics for Genomics with Dynamics Characterization of an Expanded KRAS Mutational Landscape": Supplmental Information

### SUPPLEMENTAL INFORMATION

Table S1: The number of experimentally solved KRAS structures, per mutational and ligand status.

| Mutation | Nucleotide | Fully Resolved (1-169) | In complex | Not Fully Resolved | In complex |
|---|---|---|---|---|---|
| WT | GTP | 17 | 6 | 10 | 9 |
| | GDP | 22 | 6 | 12 | 3 |
| | apo | - | - | 1 | 1 |
| G12C | GTP | - | - | 1 | - |
| | GDP | 28 | - | 15 | - |
| | apo | - | - | 5 | 5 |
| G12D | GTP | 12 | 1 | 3 | - |
| | GDP | 2 | - | 6 | 2 |
| | apo | - | - | - | - |
| Q61H | GTP | 7 | - | 1 | - |
| | GDP | - | - | - | - |
| | apo | - | - | - | - |
| G12V | GTP | 1 | 1 | 4 | - |
| | GDP | 5 | - | 2 | - |
| | apo | - | - | - | - |
| G13D | GTP | 3 | 3 | 1 | - |
| | GDP | 2 | - | - | - |
| | apo | - | - | - | - |
| Q61R | GTP | 2 | 2 | - | - |
| | GDP | - | - | - | - |
| | apo | - | - | - | - |
| P34R | GTP | 2 | - | 1 | - |
| | GDP | - | - | 1 | - |
| | apo | - | - | - | - |
| G12A | GTP | - | - | 3 | - |
| | GDP | 1 | - | 1 | - |
| | apo | - | - | - | - |
| G12R | GTP | - | - | 1 | - |
| | GDP | 1 | - | - | - |
| | apo | - | - | - | - |
| V14I | GTP | - | - | - | - |
| | GDP | 1 | - | 1 | - |
| | apo | - | - | - | - |
| A146T | GTP | - | - | - | - |
| | GDP | 1 | - | - | - |
| | apo | - | - | - | - |
| A49G | GTP | 1 | - | - | - |

| | | | | | |
|---|---|---|---|---|---|
| | GDP | - | - | - | - |
| | apo | - | - | - | - |
| A59G | GTP | - | - | - | - |
| | GDP | 1 | - | - | - |
| | apo | - | - | - | - |
| D33E | GTP | - | - | - | - |
| | GDP | 1 | - | - | - |
| | apo | - | - | - | - |
| Q61L | GTP | - | - | 1 | - |
| | GDP | - | - | 1 | - |
| | apo | - | - | - | - |
| Q61A | GTP | - | - | - | - |
| | GDP | 1 | - | - | - |
| | apo | - | - | - | - |
| M72C | GTP | - | - | - | - |
| | GDP | 1 | - | - | - |
| | apo | - | - | - | - |

Table S2: Complete list of scores on a per variants basis (Additional File)

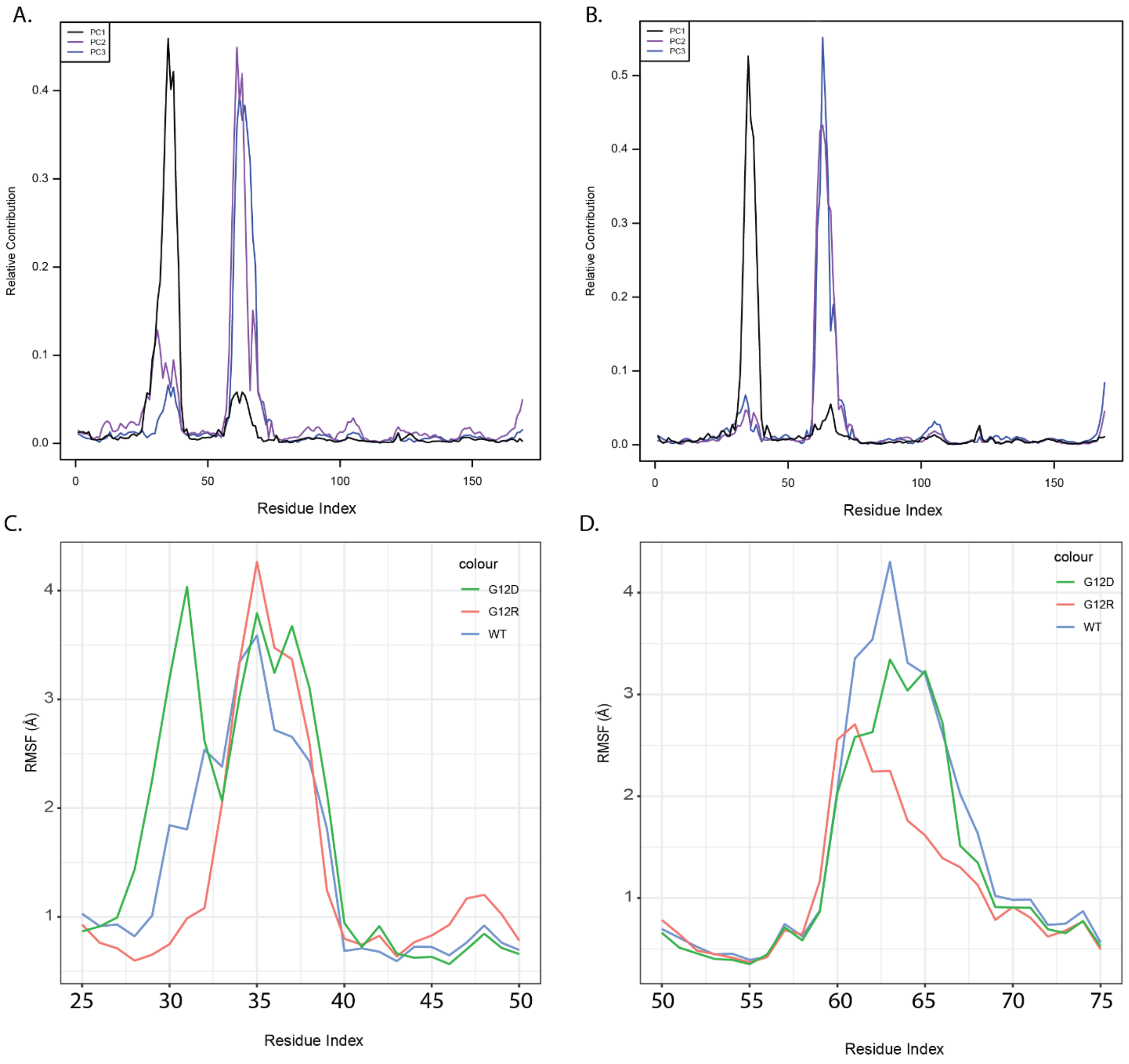


Figure S1: PCA relative contributions for A) GTP and B) GDP simulated coordinate space. GDP simulation RMSF of C) Switch 1 and D) Switch 2 for WT, G12R, and G12D.

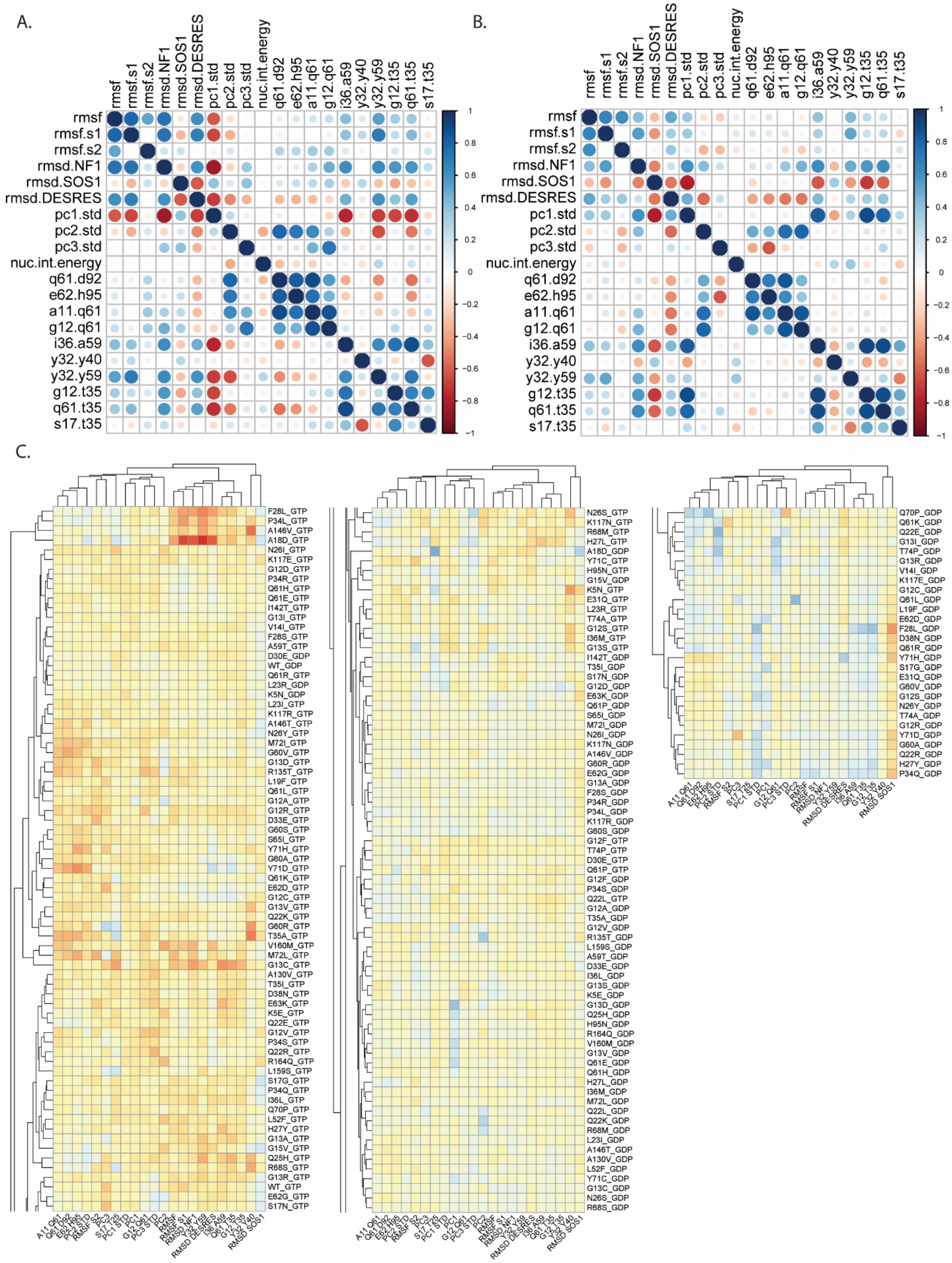
A.
B.
C.
rmsf
rmsf.s1
rmsf.s2
rmsd.NF1
rmsd.SOS1
rmsd.DESRES
pc1.std
pc2.std
pc3.std
nuc.int.energy
q61.d92
e62.h95
a11.q61
g12.q61
i36.a59
y32.y40
y32.y59
g12.t35
q61.t35
s17.t35
1
0.8
0.6
0.4
0.2
0
−0.2
−0.4
−0.6
−0.8
−1
F28L_GTP
P34L_GTP
A146V_GTP
A18D_GTP
N26I_GTP
K117E_GTP
G12D_GTP
P34R_GTP
Q61H_GTP
Q61E_GTP
I142T_GTP
G13I_GTP
V14I_GTP
F28S_GTP
A59T_GTP
D30E_GDP
WT_GDP
Q61R_GTP
L23R_GDP
K5N_GDP
L23I_GTP
K117R_GTP
A146T_GTP
N26Y_GTP
M72I_GTP
G60V_GTP
G13D_GTP
R135T_GTP
L19F_GTP
Q61L_GTP
G12A_GTP
G12R_GTP
D33E_GTP
G60S_GTP
S65I_GTP
Y71H_GTP
G60A_GTP
Y71D_GTP
Q61K_GTP
E62D_GTP
G12C_GTP
G13V_GTP
Q22K_GTP
G60R_GTP
T35A_GTP
V160M_GTP
M72L_GTP
G13C_GTP
A130V_GTP
T35I_GTP
D38N_GTP
E63K_GTP
K5E_GTP
Q22E_GTP
G12V_GTP
P34S_GTP
Q22R_GTP
R164Q_GTP
L159S_GTP
S17G_GTP
P34Q_GTP
I36L_GTP
Q70P_GTP
L52F_GTP
H27Y_GTP
G13A_GTP
G15V_GTP
Q25H_GTP
R68S_GTP
G13R_GTP
WT_GTP
E62G_GTP
S17N_GTP
N26S_GTP
K117N_GTP
R68M_GTP
H27L_GTP
A18D_GDP
Y71C_GTP
H95N_GTP
G15V_GDP
K5N_GTP
E31Q_GTP
L23R_GTP
T74A_GTP
G12S_GTP
I36M_GTP
G13S_GTP
I142T_GDP
T35I_GDP
S17N_GDP
G12D_GDP
E63K_GDP
Q61P_GDP
S65I_GDP
M72I_GDP
N26I_GDP
K117N_GDP
A146V_GDP
G60R_GDP
E62G_GDP
G13A_GDP
F28S_GDP
P34R_GDP
P34L_GDP
K117R_GDP
G60S_GDP
G12F_GTP
T74P_GTP
D30E_GTP
Q61P_GTP
G12F_GDP
P34S_GDP
Q22L_GTP
G12A_GDP
T35A_GDP
G12V_GDP
R135T_GDP
L159S_GDP
A59T_GDP
D33E_GDP
I36L_GDP
G13S_GDP
K5E_GDP
G13D_GDP
Q25H_GDP
H95N_GDP
R164Q_GDP
V160M_GDP
G13V_GDP
Q61E_GDP
Q61H_GDP
H27L_GDP
I36M_GDP
M72L_GDP
Q22L_GDP
Q22K_GDP
R68M_GDP
L23I_GDP
A146T_GDP
A130V_GDP
L52F_GDP
Y71C_GDP
G13C_GDP
N26S_GDP
R68S_GDP
Q70P_GDP
Q61K_GDP
Q22E_GDP
G13I_GDP
T74P_GDP
G13R_GDP
V14I_GDP
K117E_GDP
G12C_GDP
Q61L_GDP
L19F_GDP
E62D_GDP
F28L_GDP
D38N_GDP
Q61R_GDP
Y71H_GDP
S17G_GDP
E31Q_GDP
G60V_GDP
G12S_GDP
N26Y_GDP
T74A_GDP
G12R_GDP
Y71D_GDP
G60A_GDP
Q22R_GDP
H27Y_GDP
P34Q_GDP
A11 Q61
Q61 D92
E62 H95
PC2 STD
RMSF S2
PC3
S17 T35
PC1 STD
PC1
G12 Q61
PC3 STD
PC2
RMSF
RMSF S1
RMSD NF1
Y32 Y59
RMSD DESRES
I36 A59
Q61 T35
G12 T35
Y32 Y40
RMSD SOS1

Figure S2: Summary of interconnectedness between MD metrics. Correlation matrix between all MD metrics for A) GDP and B) GTP simulations. C) Heatmap of all score values on a per variant basis. Red values indicate higher score values, while blue values indicate of lower score values. Scores are clustered via k-means on the x-axis and variants are clustered via k-means on the y-axis.

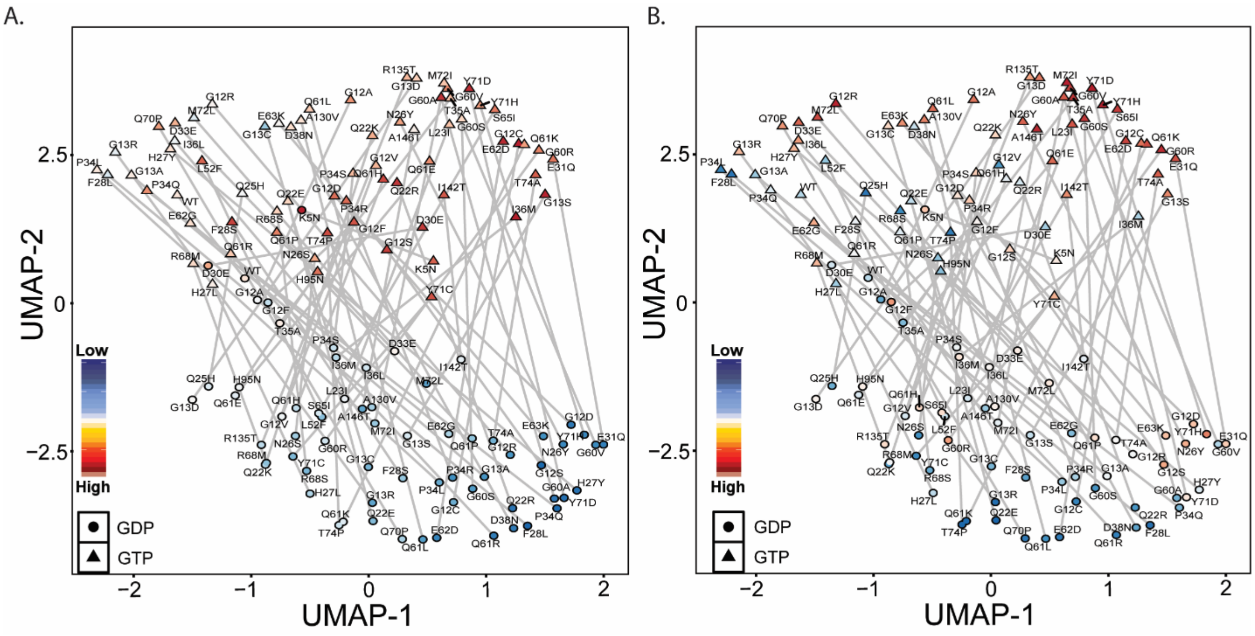


Figure S3: Observing PC Standard Scores over UMAP exemplifies the relative importance of features.

A) UMAP dimensionality reduction calculated using all MD metrics discussed in this study. Each variant is colored according to the PC1 standard score and using the same format as Figure 4A. and B) according to the PC2 standard score.
